# Divergent thermal challenges elicit convergent stress signatures in aposymbiotic *Astrangia poculata*

**DOI:** 10.1101/2020.01.25.919399

**Authors:** DM Wuitchik, A Almanzar, BE Benson, SA Brennan, JD Chavez, MB Liesegang, JL Reavis, CL Reyes, MK Schniedewind, IF Trumble, SW Davies

## Abstract

Anthropogenic climate change threatens corals globally and both high and low temperatures are known to induce coral bleaching. However, coral stress responses across wide thermal breadths are rarely explored. In addition, it is difficult to disentangle the role of symbiosis on the stress response of obligately symbiotic coral hosts. Here, we leverage aposymbiotic colonies of the facultatively symbiotic coral, *Astrangia poculata*, which lives naturally with and without its algal symbiont, to examine how broad thermal challenges influence coral hosts. *A. poculata* were collected from their northern range limit and thermally challenged in two independent 16-day common garden experiments (heat and cold challenge) and behavioral responses to food stimuli and genome-wide gene expression profiling (TagSeq) were performed. Both thermal challenges elicited significant reductions in polyp extension. However, five times as many genes were differentially expressed under cold challenge compared to heat challenge. Despite more genes responding to cold challenge, there was significant overlap in which genes were differentially expressed across thermal challenges. These convergently responding genes (CRGs) were associated with downregulation of motor functions and nematocysts while others were consistent with stress responses previously identified in tropical corals. The fact that these responses were observed in aposymbiotic colonies highlights that many genes previously implicated in stress responses in symbiotic species may simply represent the coral’s stress response in or out of symbiosis.

## Introduction

Temperature is an important factor in determining species distribution patterns in ectothermic organisms (Angilleta 2009). As sea surface temperatures continue to rise, understanding how these changes will affect species distributions demands a broad understanding of organisms’ physiological sensitivities to temperature across their native range. There is overwhelming evidence that temperature increases associated with anthropogenic climate change are having widespread ecological consequences on marine species distributions (Hoegh-Guldberg *et al*. 2008; Pinsky *et al*. 2019). Coral reefs are particularly sensitive to these thermal changes, which have been implicated in widespread reef declines (Hughes *et al*. 2017). Temperature anomalies are the primary driver of the breakdown in the obligate symbiotic relationship between tropical corals and their endosymbiotic algae (family Symbiodiniaceae; LaJeunesse *et al*. 2018). This breakdown results in the expulsion of algae from coral host tissue in a process known as coral bleaching (Gates *et al*. 1992; Venn *et al*. 2008). Because symbiotic algae translocate carbon sugars to the coral host, losing these symbionts results in significant energy loss and many corals are unable to survive extended periods in a bleached state (Weis 2008).

The majority of research on coral bleaching has focused on responses to elevated temperatures in tropical reef-building corals (Cziesielski *et al*. 2019). However, tropical corals can bleach in response to a variety of stressors, including high nutrients (Wiedenmann *et al*. 2013), ocean acidification (Anthony *et al*. 2008), pathogens (Ben-Haim & Rosenberg 2002), low salinity (Goreau 1964), chemical exposures (Cervino *et al*. 2003), and cold stress (Saxby *et al*. 2003). Coral responses to cold stress remain understudied, even though these events can have substantial impacts on reefs. For example, a cold-water bleaching event in 2010 decimated inshore coral populations along the Florida reef tract (Lirman *et al*. 2011), and cold water has caused bleaching on the Great Barrier Reef (Hoegh-Guldberg & Fine 2004). While the main effect of climate change on marine systems is a net increase in mean global sea surface temperatures, these cold thermal challenges may be exacerbated by the pace of Arctic warming (twice as fast as the global average), which may influence upper-level atmospheric activity and storm tracks resulting in more frequent extreme cold outbreaks at northern mid-latitudes (Cohen *et al*. 2014). These cold extremes are therefore relevant thermal challenges to subtropical and temperate coral species.

One way to monitor responses to stress is to characterize changes in gene expression profiles, which provides a snapshot into the physiological state of an organism and offers insights into the biological processes, molecular functions, and cellular components that corals engage to tolerate various stressors. Modern transcriptomics have demonstrated that corals mount dynamic responses to pollutants (Gust *et al*. 2014; Ruiz-Ramos *et al*. 2017), pH (Moya *et al*. 2012; Davies *et al*. 2016) and bacterial challenges (Fuess *et al*. 2017; Wright *et al*. 2017) and considerable efforts have been made to understand how corals respond to heat challenges (for review see Cziesielski *et al*. 2019). Interestingly, similar patterns of gene expression emerge from these different stressors. Barshis *et al*. (2013) demonstrates that corals exhibit a widespread stress response across thousands of genes, and this environmental stress response (ESR) is consistent with the conserved response to diverse environmental stressors in yeast (Gasch *et al*. 2000). A meta-analysis comparing the transcriptomic responses of coral from the genus *Acropora* to various stressors found these coral exhibit a stereotyped ESR (Dixon *et al*. 2020). There, it was found that there is consistent upregulation of genes involved in cell death, response to reactive oxygen species, NF-kB signaling, immune response, protein folding, and protein degradation to a variety of acute stress exposures. This research highlights that testing a single stressor cannot elucidate whether genes being expressed are unique to the stressor or emerge from a more generalized ESR.

Most work exploring the stress responses of corals have focused on tropical reef-building corals that live in oligotrophic waters and cannot survive long-term without their algal symbionts. Because energy deprivation in coral hosts results from any mechanism of symbiont loss (Baena-González & Sheen 2008), uncoupling a thermal stress response from an energy deprivation response is challenging. Furthermore, given that many tropical corals exhibit an obligate symbiotic relationship, it is difficult to disentangle the host’s stress response to extreme temperatures from the host’s response to stress-induced algal by-products (i.e. reactive oxygen species (ROS); McGinty *et al*. 2012) and the resulting energy deprivation from dysbiosis. However, there are several species of subtropical and temperate reef-building corals that exhibit facultative symbioses and offer promising avenues to better understand stress responses.

The Northern Star Coral *(Astrangia poculata)* exhibits a facultatively symbiotic relationship with *Breviolum psygmophilum* (LaJeunesse *et al*. 2012) and can be found in sympatry in varying symbiotic states that are visually distinguishable by colour. Symbiotic colonies appear brown due to high densities of *B. psygmophilum*, and much like a bleached coral, some *A. poculata* appear white (Figure 1C) due to very low algal densities (Dimond & Carrington 2007). This white phenotype is commonly referred to as “aposymbiotic” (Grace 2017; Sharp *et al*. 2017; Burmester *et al*. 2018) due to the paucity of algal symbionts. Unlike obligate symbiotic corals, *A. poculata* can thrive in its aposymbiotic state relying only on heterotrophy (Dimond & Carrington 2007). Additionally, *A. poculata* experiences large seasonal variation in temperature at its northern range, making these populations ideal models for investigating how corals might withstand wide thermal challenges. Taken together, aposymbiotic *A. poculata* provide a unique opportunity to disentangle how broad thermal challenges influence the coral host in isolation from its algal symbiont. Here, we present two thermal challenge experiments that independently assess the behavioural and molecular responses of aposymbiotic *A. poculata* to divergent thermal challenges.

**Figure 1.**
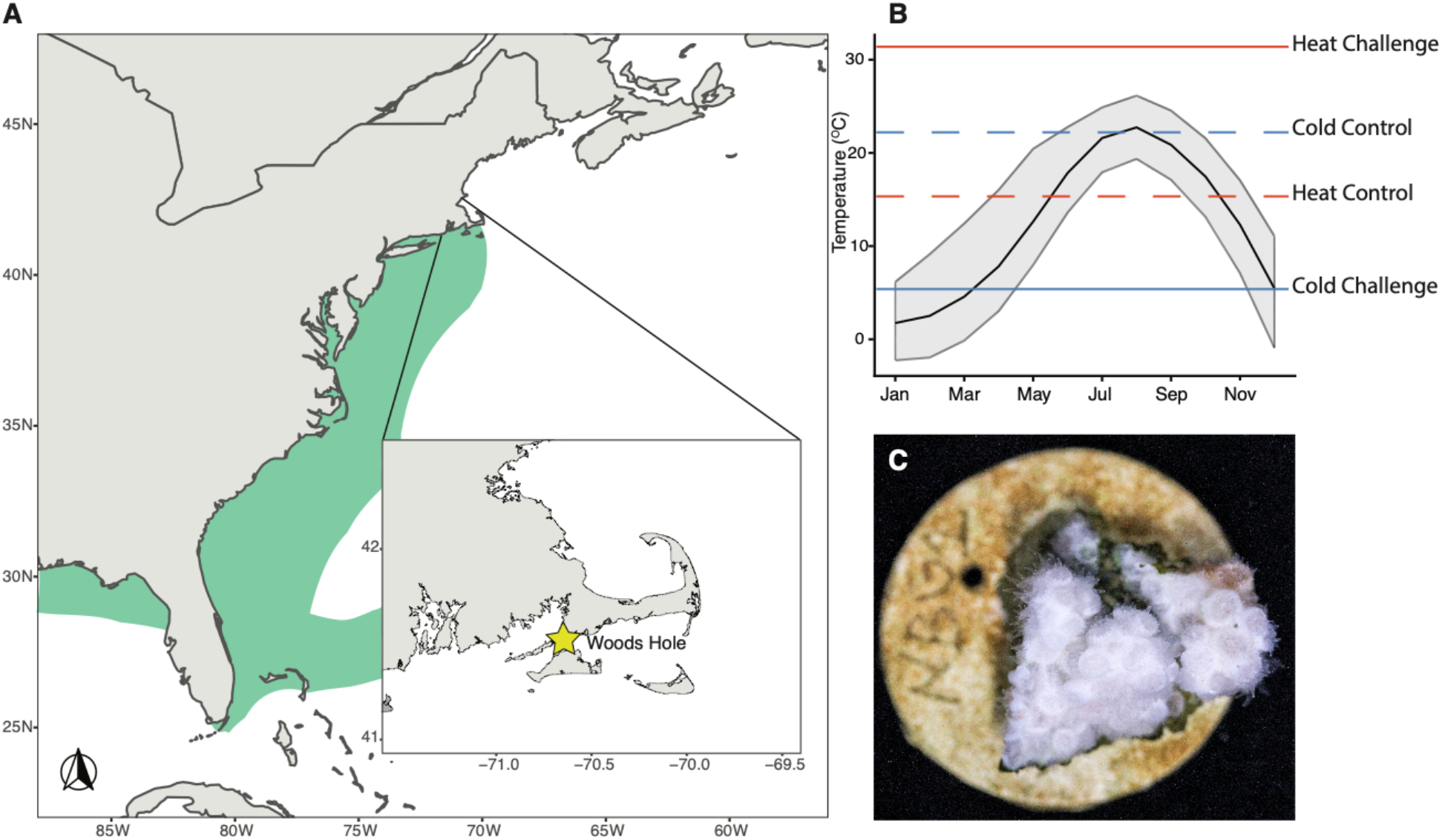
**A)** Map of the eastern seaboard of the United States with the *Astrangia poculata* range in green. Inset shows the Woods Hole collection site denoted with a yellow star (distributions based on Thornhill *et al*. 2008). **B)** Seasonal temperature profile at Woods Hole averaged over ten years (2008-2018). The black solid line indicates mean monthly temperatures with mean monthly maximum and minimum temperatures in grey. Temperatures (°C) of thermal challenge experimental controls (dashed lines) and treatments (solid lines) are superimposed with cold challenge treatments in blue and heat challenge treatments in red. Seasonal temperatures were obtained from the National Oceanic and Atmospheric Administration weather buoy #BZBM3. **C)** Picture of an aposymbiotic *A. poculata* colony fragment.

## Methods

### Thermal challenge common garden experiments

Eighteen unique aposymbiotic colonies of *Astrangia poculata* were collected in Woods Hole, Massachusetts (41.54N, 70.64W; Figure 1A) in October, 2017 and transported to the Marine Invertebrate Research Facility at Boston University. Colonies were acclimated at 16°C for three weeks. On November 17, 2017, colonies were fragmented, each coral nubbin was assigned a unique ID and glued to a labelled dish (Figure 1C). Nubbins were allowed to recover from fragmentation and further acclimated at 16°C under a 12 L:12 D photoperiod with light levels ranging from 6-12 μmol m^-2^s^-1^ and fed *Artemia spp*. nauplii daily for 24 days.

### Thermal challenge I: cold challenge experiment

Nine unique aposymbiotic colonies were assigned to the cold challenge experiment (Table 1). At least one nubbin from each colony was represented in one of three replicate tanks assigned to control conditions (maintained at 22°C) and one of three replicate tanks assigned to the cold challenge treatment (incrementally lowered from 23°C by approximately 1°C/day to a final temperature of 6°C; Figure 2A). When additional fragments remained from a colony, they were randomly stratified into different tanks (n=43 nubbins total). Several aspects of this experimental design are noteworthy. The first is that our control treatment (22°C) was 6°C higher than the coral acclimation temperature, which may have caused an initial stress response in the first few days. The second aspect is that 6°C is warmer than the minimum temperature *A. poculata* experience within their seasonal averages (Figure 1B); however, achieving lower temperatures was limited by the capacity of our aquarium chillers. In addition, these colonies were collected in October so this thermal minimum and the rate at which this minimum was achieved likely represents a considerable thermal challenge for these corals.

**Table 1.** Summary of distribution of coral genotypes among treatments for the A) cold challenge and B) heat challenge experiment. Numbers in cells represent the number of fragments of each genotype in each treatment; numbers in parentheses represent the number of fragments that were successfully sequenced via TagSeq.

| A) Cold Challenge Experiment |  |  | B) Heat Challenge Experiment |  |  |
| --- | --- | --- | --- | --- | --- |
| Genotype | Control (TagSeq) | Challenge (TagSeq) | Genotype | Control (TagSeq) | Challenge (TagSeq) |
| D | 1 (1) | 1 (0) | A | 1 (1) | 3 (1) |
| E | 3 (2) | 3 (1) | B | 3 (2) | 3 (2) |
| F | 3 (2) | 3 (2) | C | 3 (2) | 3 (0) |
| G | 2 (0) | 2 (1) | D | 3 (2) | 3 (1) |
| H | 3 (2) | 1 (1) | F | 1 (1) | 1 (0) |
| I | 2 (2) | 2 (0) | K | 2 (0) | 2 (1) |
| J | 3 (2) | 3 (2) | N | 3 (0) | 1 (1) |
| P | 1 (1) | 3 (2) | O | 2 (1) | 2 (2) |
| T | 3 (2) | 3 (3) | Q | 3 (3) | 3 (2) |
| Total | 21 (14) | 21 (12) |  | 21 (12) | 21 (10) |

**Figure 2.**
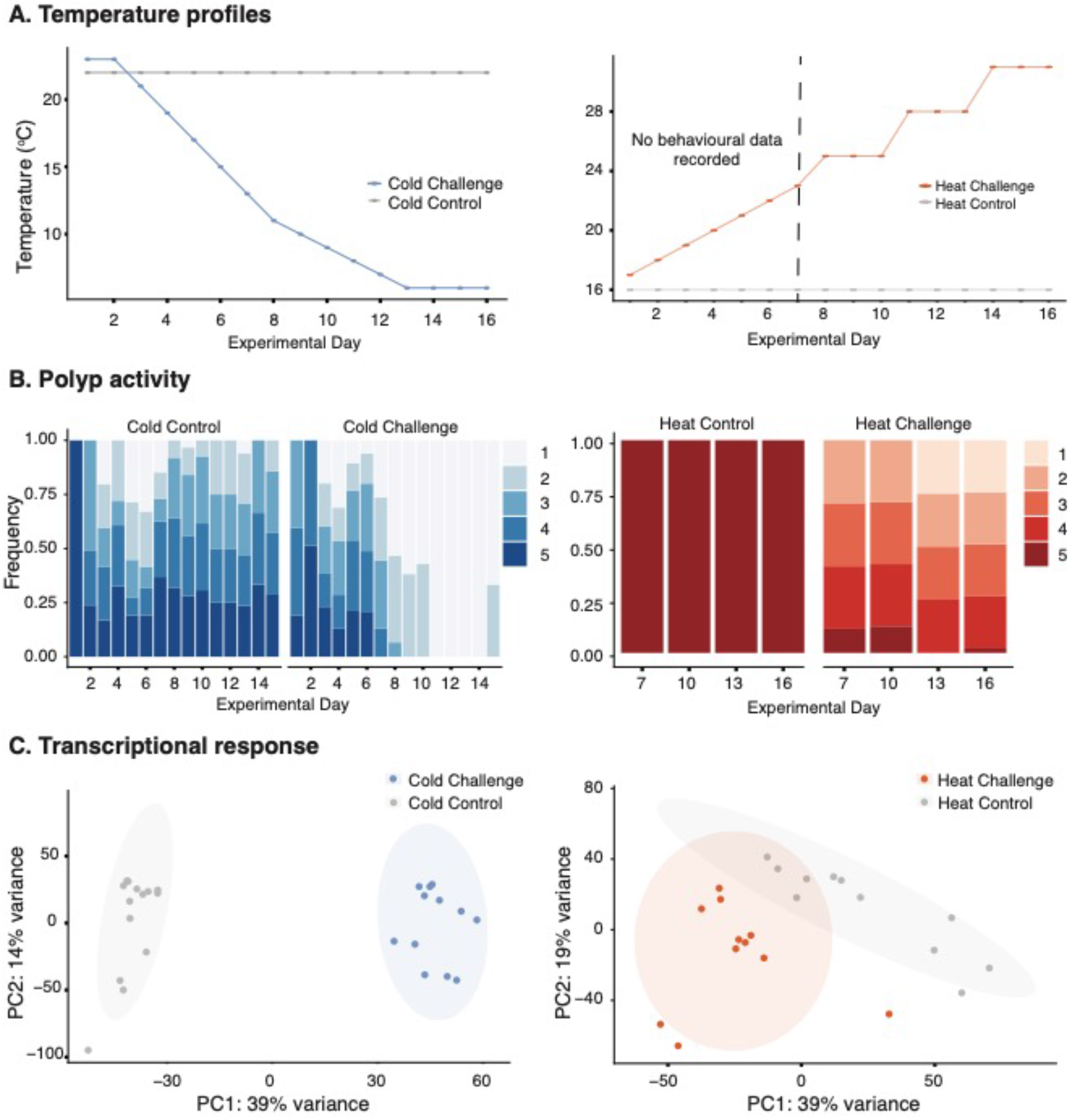
Thermal challenge experiments on *Astrangia poculata*. Left: cold challenge, Right: heat challenge. A) 16-day temperature ramp. B) Polyp activity scored based on the proportion of polyps extended per fragment (1 = 0%, 2 = 25%, 3 = 50%, 4 = 75%, 5 = 100%) in response to food stimuli across the 16-day experiments. Note that behavioral data collection in the heat challenge experiment did not commence until day 7. C) Principal component analysis of overall gene expression of samples under control and thermal challenge at day 16. Percentages represent the total variance explained by each axis and shaded areas are 95% confidence ellipses. P-value indicates significance of treatment using a permutational multivariate analysis of variance.

### Thermal challenge II: heat challenge experiment

An independent set of nine unique aposymbiotic *A. poculata* colonies were fragmented and at least one nubbin from each colony was assigned to each treatment. There were three tank replicates for control conditions (maintained at 16°C) and three replicate tanks for the heat challenge treatment. At least one nubbin of each colony was assigned to each treatment, and when additional colony fragments remained, they were randomly stratified into different tanks (Table 1). At the beginning of the 16 day heat challenge experiment, all tanks were maintained at 16°C. Heat challenge tanks were ramped from 16°C to 23°C over 6 days (approximately 1°C/day) but no phenotypic data were recorded during this time. Phenotype observations were conducted on days 7-16 during which heat challenge tanks were incrementally ramped 2-3°C in one day followed by a 2-day recovery period. This ramping protocol continued until 31°C was achieved (Figure 2A). It is worth noting several aspects of this experimental design: the final heat challenge temperature was well above the maximum temperature these corals experience at their source location (Figure 1B) and the heat challenge experiment was conducted independently from the cold challenge experiment described above (Figure 2A).

### Coral polyp behaviour in response to food stimulus

In the cold challenge experiment, corals were fed daily and feeding behaviours were recorded 30 minutes after feeding. In contrast, in the heat challenge experiment, phenotypic measurements were not conducted in the first 6 days. Heat challenge phenotypic measurements began on day 7 and continued after corals were offered food every third day for the duration of the experiment (16 days). Coral polyp behaviour in response to food stimulus was quantified by the total coral surface area that had observable polyp extension relative to retracted polyps. This score was on a scale of 1 to 5 based on the estimated percentage of active polyps within a fragment (1 = 0%, 2 = 25%, 3 = 50%, 4 = 75%, 5 = 100%, similar to Burmester *et al*. 2018) and the same researcher conducted all behavioural assays within each thermal challenge experiment to limit observer biases. An ordered logistic regression was performed to establish if temperature influenced polyp extension rates using the *polr* function as part of the *MASS* package (version 7.3-51.1; Venables & Ripley 2002) in R.

### Global gene expression profiling

Upon reaching maximum thermal differences between challenge and control treatments in both experiments (Day 16), several white polyps from all colonies were sampled using sterilized bone cutters, immediately placed in 200 proof ethanol and stored at −80°C. Total RNA was extracted using an RNAqueous kit (Ambion by LifeTechnologies) following the manufacturer’s recommendations. An additional step was implemented using 0.5 mm glass beads (BioSpec), which were added to the vial of lysis buffer and samples were homogenized using a bead beater for 1 min. RNA quantity and integrity were determined using a DeNovix DS-11+ spectrophotometer and ribosomal RNA bands were confirmed on 1% agarose gels. Trace DNA contamination was removed using a DNase 1 (Ambion) digestion at 37°C for 45 minutes. Libraries were created from 1500 ng of total RNA (following Meyer *et al*. 2011) and adapted for Illumina Hi-Seq sequencing (Dixon *et al*. 2015; Lohman *et al*. 2016). In brief, RNA was heat-sheared and transcribed into first-strand cDNA using a template-switching oligo and SMARTScribe reverse transcriptase (Clontech). cDNA was then PCR-amplified, individual libraries were normalized, and Illumina barcodes were incorporated using a secondary PCR. Samples were pooled and size-selected prior to sequencing on Illumina Hiseq 2500 single-end (SE) 50 basepair (bp) at Tufts University Core Facility (TUCF). Due to insufficient RNA yield, some samples were not successfully represented in library preparations. Of the 42 samples within each of the cold and heat challenge experiments, 26 and 22 libraries were prepared, respectively (Table 1).

### Transcriptome assembly and gene expression analyses

Illumina TruSeq adapters and poly-A tails were first removed using the *FASTX-Toolkit* (v 0.0.14, Hannon, G.J. (2010) FASTX-Toolkit. http://hannonlab.cshl.edu/fastx_toolkit.) and resulting sequences that were less than 20 bp in length were removed. In addition, only those sequences with > 90% of bases having a quality score > 20 were retained. PCR duplicates were removed and resulting quality-filtered reads were concatenated and used to assemble a novel transcriptome using Trinity (Grabherr *et al*. 2013). Contigs were then annotated using BLAST (Altschul *et al*. 1990) searches against UniProt and Swiss-Prot NCBI NR protein databases. This newly assembled transcriptome along with its annotation files are included as Supplementary Files 1-4 (1: transcriptome fasta file, 2: seq2iso file, 3: iso2gene, 4: iso2GO and are also available at http://sites.bu.edu/davieslab/data-code/).

Quality-filtered reads were then mapped to the newly assembled transcriptome using *Bowtie2* (Langmead & Salzberg 2012). There were an average 520,662 mapped reads across both experiments with mapping efficiencies ranging from 36%-57% (Supplementary File 5). Raw count files for each experiment are available in Supplemental Files 6 (cold challenge) and 7 (heat challenge). Data from each challenge experiment were analyzed independently. First, data were tested for outliers using *arrayQualityMetrics* as part of DESeq (Anders & Huber 2010) and no outliers were detected for either experiment. DESeq2 (Love *et al*. 2014) was then used to identify differentially expressed genes (DEGs, Supplemental Files 8-9) associated with cold and heat thermal challenge relative to their respective controls using a Wald’s test. P-values were adjusted for multiple testing using the Benjamini and Hochberg method (FDR < 0.05; Benjamini & Hochberg 1995). Lastly, expression data for each experiment were *r-log* transformed and these data were used as input for a principal component analysis. A permutational multivariate analysis of variance was then used to determine if overall gene expression patterns between thermal challenge treatments differed significantly from their controls using the *adonis* function in vegan v2.5-4 (Oksanen *et al*. 2019).

Gene ontology (GO) enrichment analyses were performed using adaptive clustering of GO categories and Mann–Whitney U tests (GO-MWU) based on the ranking of signed log p-values (Voolstra *et al*. 2011), which is particularly suitable for non-model organisms (Dixon *et al*. 2015). Results were visualized in dendrograms tracing the level of gene sharing between significant categories and direction of change in treatment temperatures compared to their respective controls.

### Testing for a convergent response to thermal challenge

Lists of DEGs (FDR < 0.05) between the two thermal challenge experiments were compared and visualized using a Venn Diagram; and, significant enrichment of genes at the intersection between experiments was tested for using a hypergeometric test. The DEGs at the intersection between experiments (common DEGs) were visualized based on log2 fold change for each experiment; and, the most highly up- and downregulated genes were highlighted and defined as convergently responding genes (CRGs). GO categories that were independently identified as enriched (FDR < 0.05) in both experiments were visualized by their respective delta-ranks of enrichment to demonstrate the conservation of GO function across the thermal challenges (for details, see Dixon *et al*. 2015).

## Results

### Astrangia poculata *response to cold challenge*

Although behavioural responses of *A. poculata* to a food stimulus under control conditions varied, nearly all colonies exhibited some polyp extension (Figure 2B). This contrasts with behaviours observed under cold challenge, where rapid declines in polyp activity were observed by day eight (12°C) and most polyps remained inactive as cooler temperatures were reached (10°C - 6°C, Figure 2B). Overall, *A. poculata* polyp activity was significantly reduced under cold challenge (p < 0.01). *A. poculata* gene expression was also significantly influenced by cold challenge: a strong treatment effect on overall gene expression was observed (*Adonis* p_treatment_ < 0.001, Figure 2C), with cold challenge resulting in 5318 (40%) DEGs (FDR < 0.05; 2244 (17%) upregulated; 1, 3074 (23%) downregulated). Many GO terms were also enriched between cold challenge and control conditions (FDR < 0.10; CC = 77, MF =50, BP = 78; Figure 3). Of these, notable GO terms include: *myosin complex* (GO:0016459), *proteasome core complex* (GO:0005839), *translation regulator activity, nucleic acid binding* (GO:0008135; GO:0090079), *extracellular matrix structural constituent* (GO:0005201), *muscle system process* (GO: 0006936; GO:0003012) and *proteolysis* (GO:0006508).

**Figure 3.**
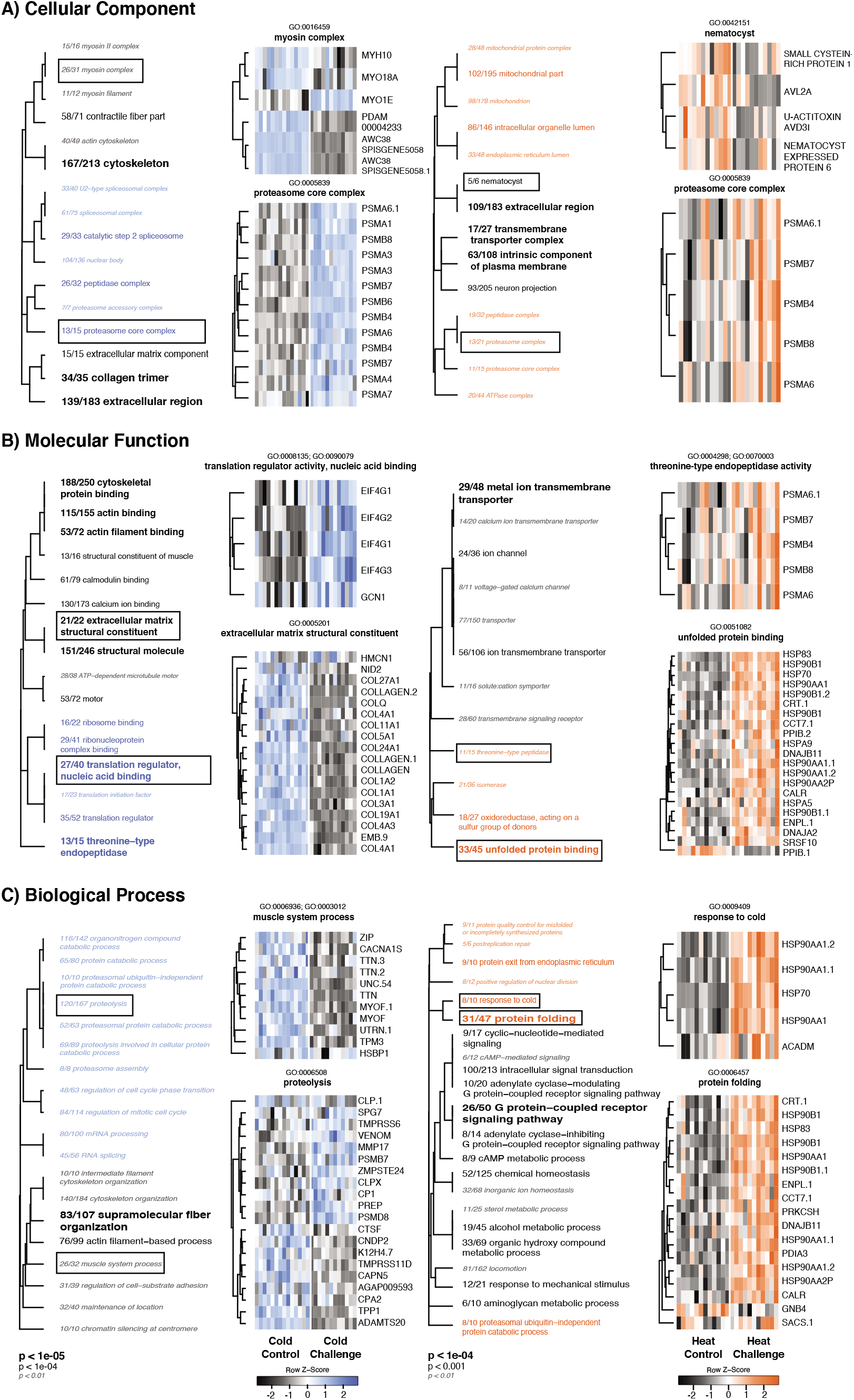
Gene ontology (GO) enrichment under thermal challenges: Left: cold challenge, Right: heat challenge. Enriched GO terms of A) Cellular Components B) Molecular Functions, and C) Biological Processes were determined via Mann-Whitney U tests. Font size and boldness of text corresponds to p-values with colour designating directionality of enrichment (blue: cold challenge, red: heat challenge, black: controls). GO terms are clustered based on the number of shared genes between categories. Hierarchical clustered heatmaps were generated from annotated DEGs with a highlighted GO term (black box) and each row was labelled with its gene symbol. Colors denote magnitude of response (blue: upregulated in cold challenge, red: upregulated in heat challenge) through z-score of the difference in expression levels from that of mean expression for each gene.

### Astrangia poculata *response to heat challenge*

Behavioural responses of *A. poculata* to a food stimulus under control conditions were stable and coral polyps remained fully extended throughout the experiment (Day 7 - 14; Figure 2B). This contrasts with behavioural responses under heat challenge, where corals exhibited less polyp activity in response to food stimulus as temperatures increased. By the end of the experiment (day 16), only one colony under heat challenge was observed to have 100% polyp extension and half of the colonies had less than 25% of their polyps extended (Figure 2B). Overall, *A. poculata* polyp activity was significantly reduced under heat challenge (p < 0.01). *A. poculata* gene expression was also significantly influenced by heat challenge: a significant effect of treatment on overall gene expression was observed *(Adonis* p_treatment_< 0.001, Figure 2C) with 1,054 (7.9%) DEGs (FDR < 0.05; 410 (3.1%) upregulated; 644 (4.9%) downregulated. Many GO terms were significantly enriched under heat challenge relative to control conditions (FDR < 0.10; CC = 34, MF = 47, BP = 22; Figure 3). Notable GO terms include: *nematocyst* (GO:0042151), *proteasome core complex* (GO:0005839), *threonine-type endopeptidase activity* (GO:0004298; GO:0070003), *unfolded protein binding* (GO:0051082), *protein folding* (GO:0006457) and *response to cold* (GO:0009409).

### *Convergent response repertoires to heat and cold challenge in* Astrangia poculata

Both cold and heat thermal challenges induced a reduction in polyp activity in response to food stimulus (Figure 2B). However, this reduction was more pronounced under cold challenge where nearly all polyps were retracted by day 16. In addition, five times as many genes were differentially expressed under cold challenge compared to heat challenge (Figure 4A). More than half (657 out of 1054) of DEGs in the heat challenge experiment were also differentially expressed under cold challenge, which is significantly more genes shared between experiments than would be expected by chance (hypergeometric test, p < 0.01). Genes that were highly upregulated under both thermal challenges include: *tumour necrosis receptor 3* (TRAF3), *Lon protease 2, peroxisomal* (LONP2), and *increased sodium tolerance 1* (ITS1). Genes that were highly downregulated under both thermal challenge treatments include: *DELTA-thalatoxin-AVl2a* (AVL2A)*, myosin regulatory light polypeptide 9* (MYL9), and *Protein-glucosylgalactosylhydroxylysine glucosidase* (PGGHG). GO terms consistently enriched in both experiments were also visualized using experimental delta-ranks of enrichment for each thermal challenge (FDR < 0.10; MF = 11, BP = 4, CC = 14, Figure 5A-C). These terms included *response to mechanical stimulus* (GO:0009612) and *locomotion* (GO:0040011) as well as GO terms associated with the proteasome (GO:0004298, GO:0006515, GO:0008540, GO:0022624, and GO:0005839).

**Figure 4.**
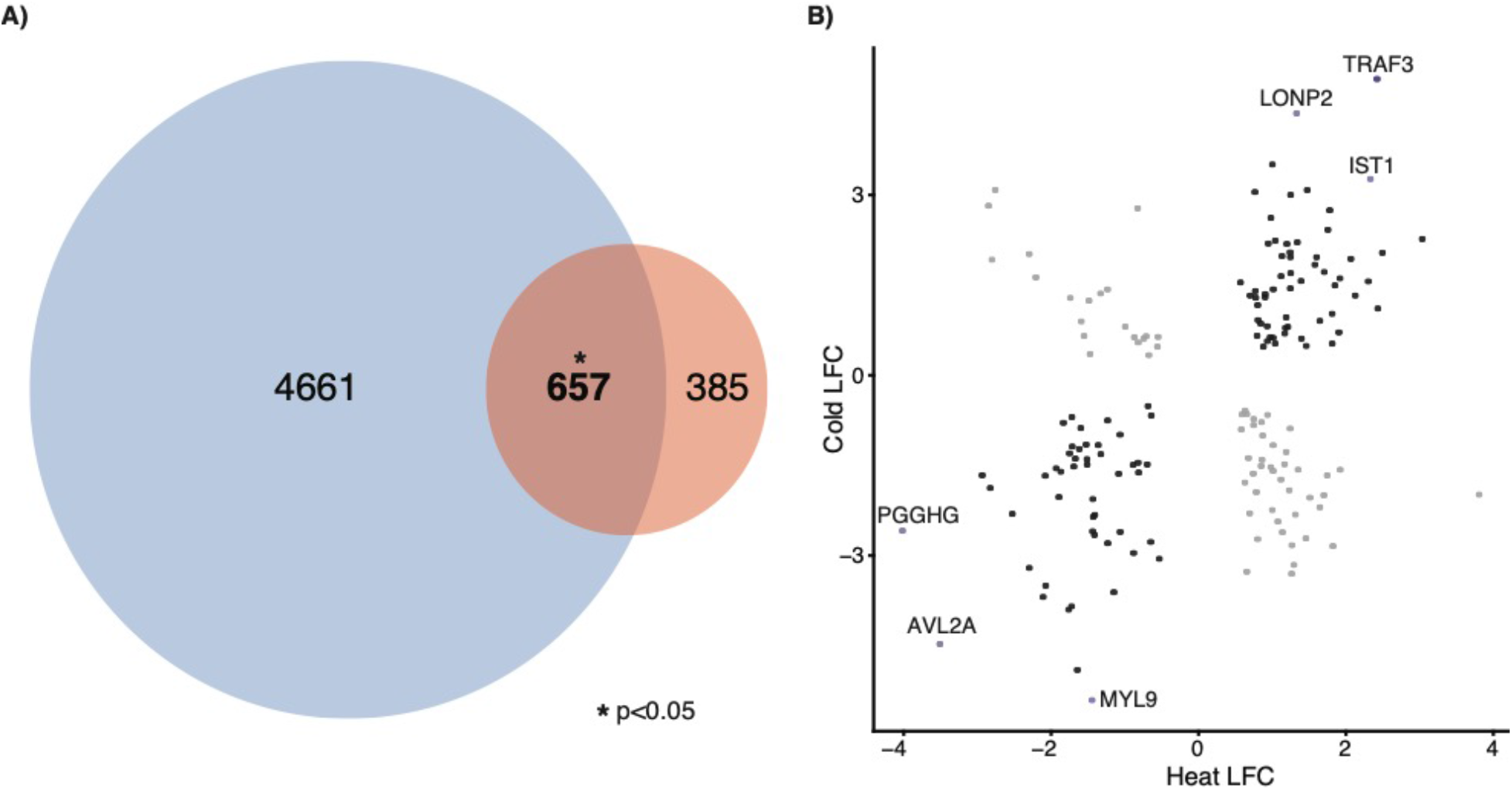
Convergent transcriptomic response of *Astrangia poculata* to thermal challenges. A) Venn diagram of differentially expressed genes shared (intersection) between cold (blue) and heat (red) challenge experiments. B) Of these 657 shared DEGs, those with annotations are visualized by their respective Log2fold change (LFC) in each experiment. Genes with consistent direction in their respective LFC are designated as convergently responsive genes (CRGs) depicted as black circles and key CRGs are highlighted in purple and labeled by gene symbol. Grey circles are divergent in response to thermal challenges.

**Figure 5.**
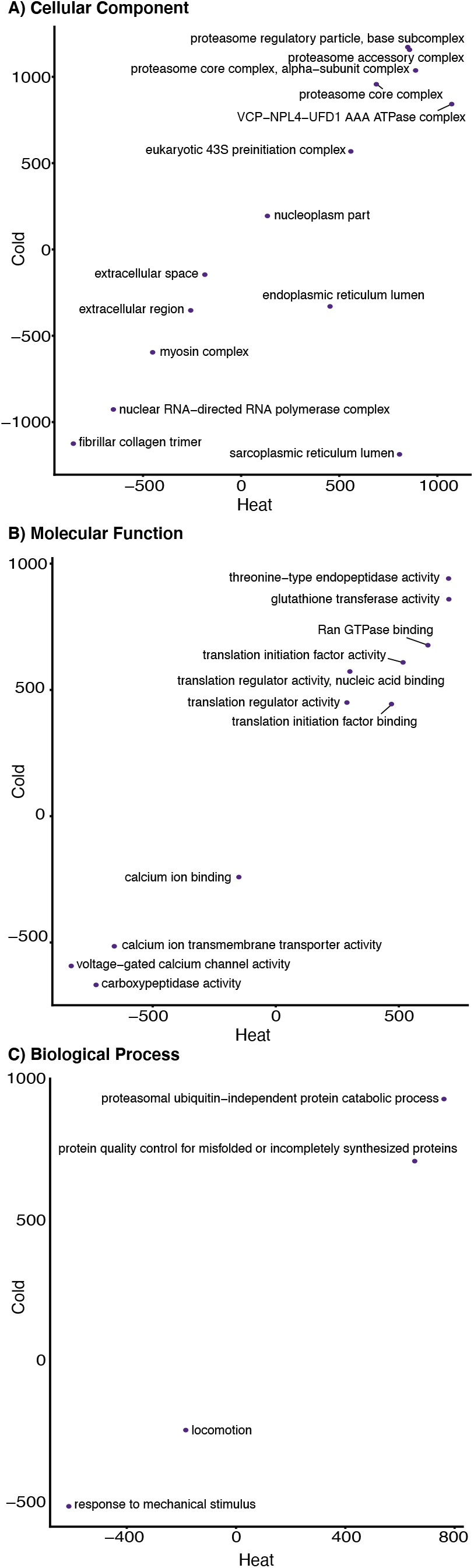
Correlation of GO delta-ranks which is the difference between mean rank of genes belonging to the GO category A) Molecular Function B) Biological Process, and C) Cellular Component, and mean rank of all other genes. Positive values indicate enrichment with up-regulated genes.

## Discussion

### *Modulation of genes associated with motor function and stress response in* Astrangia poculata *under cold challenge*

*Astrangia poculata* from Woods Hole represent the most northern range for this species with corals experiencing a wide range of temperatures throughout the year (Figure 1B). Given this temperature range, it was surprising that such strong behavioural and transcriptomic responses were observed under cold challenge (Figure 2B; 2C). This reduction in polyp activity under cold temperatures is consistent with field observations during winter months, when corals fail to respond to stimuli (e.g. quiescence, Grace 2017). The dormant polyp behaviour observed here might be interpreted as quiescence. However, very little is known about coral quiescence.

In mammalian cells, quiescent cells increase expression of certain myosin genes, notably *myosin heavy chain 10* (MYH10; Hong *et al*. 2015), which is the opposite pattern observed here under cold challenge (MYOH10 downregulated; Figure 3A). However, cold challenge did cause downregulation of other muscle responses, including *muscle system process* (MSP; GO:0006936; GO:0003012) and *myosin complex* (GO:0016459; Figure 3C), which corresponds with decreased *A. poculata* polyp activity under cold challenge. In contrast, *myosin-Ie* (MYO1E), which is an important gene for clathrin-mediated endocytosis and immunity was significantly upregulated under cold challenge. This result is consistent with previous work exploring how mice respond to cold stress (Wenzel *et al*. 2015) and myosins as a whole are often upregulated in bleached corals under heat stress (Desalvo *et al*. 2008) so it is possible that regulation of this gene is more likely related to immunity rather than muscle movement directly. The transcriptional responses match behavioral data under cold challenge, putting forth the hypothesis that reduced polyp activity under cold challenge may be regulated by downregulation of key MSP genes.

Ultimately, stress may reorganize the actin cytoskeleton in *A. poculata* under cold challenge, given the downregulation of *extracellular matrix structural constituent* (EMSC; GO:0005201). Many collagen genes are categorized within the EMSC GO category and collagen plays an integral role in forming the structure of the extracellular matrix (Kielty & Grant 2003).

Collagen genes have been previously shown to be highly reactive to a variety of stressors in corals (Traylor-Knowles 2019), which, in conjunction with these data, suggests that cold challenges lead to decreases in muscular movement and overall motor processes in aposymbiotic *A. poculata*, and potentially other corals.

Genes associated with *translation regulator activity nucleic acid binding* (GO:0008135; GO:0090079) were upregulated in *A. poculata* under cold challenge. Genes in this category are mostly composed of *eukaryotic translation initiation factor 4 gamma* (EIF4G) genes, which are consistently upregulated under a wide range of stressors, including temperature, osmotic stress and nutrient deprivation (Jones *et al*. 2013). Interestingly, EIF4G genes may play important roles for higher latitude species, like *A. poculata* here, as only northern populations of the porcelain crab *Petrolisthes cinctipes* upregulated EIF4G genes in response to cold stress (Stillman & Tagmount 2009). Given that it has already been shown that different *A. poculata* populations exhibit broadly different thermal responses (Aichelman *et al*. 2019), future work contrasting gene expression responses to stress across populations would be worthwhile.

Another GO category that demonstrated strong upregulation in *A. poculata* under cold challenge was *proteasome core complex* (PCC; GO:0005839). PCC upregulation has been previously observed in tropical corals under heat stress (Seneca & Palumbi 2015), however, this is the first to associate PCC upregulation in response to cold challenge. The majority of PCC genes are involved in the functioning of the 20S core proteasome, which is responsible for degradation of oxidized proteins (Davies 2001). Additionally, proteasomes are required for internal proteolysis of p105 into p50 to activate nuclear factor-κB (NF-κB) (Rape & Jentsch 2002). NF-κB is a key pathway in coral innate immunity and is upregulated during stress-induced bleaching in sea anemones (Mansfield *et al*. 2017).

Overall, cold challenge elicited strong effects on both behavior and transcriptomic profiles of *A. poculata* (Figure 2C); however, these patterns do not align with quiescence. Instead, these signatures are consistent with stress responses described in previous cnidarian studies and emphasize that consistent results between obligate tropical corals and aposymbiotic corals serve to highlight the host’s response to thermal challenges even in the absence of symbionts.

### *Modulation of genes associated with heterotrophy and stress response in* Astrangia poculata *under heat challenge*

Even though summer temperatures at Woods Hole over the last 10 years were much lower than temperatures achieved during the experimental heat challenge here (Figure 1B), we observed that *A. poculata* exhibited more muted behavioural and transcriptomic responses when compared to responses to cold challenge (Figure 2B; 2C). While *A. poculata* significantly reduced polyp activity in response to food stimulus under heat challenge, the majority of corals maintained some extension even at warm extremes. Unlike naturally observed polyp inactivity during winter months (Grace 2017), this is the first observation of decreased polyp activity due to warm temperatures in *A. poculata*.

Interestingly, we observed downregulation of genes associated with *nematocyst* (GO:0042151) under heat challenge, which are cnidarian stinging cells used to capture food (Holstein & Tardent 1984). Tropical corals have also been observed to reduce feeding rates under heat stress (Ferrier-Pagès *et al*. 2010). Taken together, the decreased polyp extension and downregulation of genes associated with nematocysts (Fig. 2B, 3A), suggest reduced opportunity for heterotrophy in *A. poculata*. Given that heterotrophy has been shown to mitigate coral bleaching in another facultatively symbiotic coral *(Oculina arbuscula;* Aichelman *et al*. 2016), this reduction in heterotrophy, in addition to stress associated with increased temperatures, would be interesting to explore in aposymbiotic and symbiotic colonies.

Consistent with previous work in heat challenged cnidarians, we observed upregulation of many mitochondria-related GO terms in heat challenged *A. poculata* (Figure 3). Mitochondria are fundamental in the regulation of cellular stress and have a dedicated unfolded protein response, which influences free radical detoxification and innate immunity in tropical corals (Dimos *et al*. 2019). We observed enrichment in both *protein folding* (GO:0006457) and *unfolded protein binding* (GO:0051082) under heat challenge, which is consistent with a variety of coral stress studies (i.e. Dixon et al. 2020). Genes within these GO categories are largely associated with heat shock protein production, which have been consistently implicated in coral gene expression studies (reviewed in Cziesielski *et al*. 2019) and heat stress experiments across a wide range of taxa (reviewed in Chen *et al*. 2018). Unexpectedly, we observed upregulation of *response to cold* (GO:0009409), which is a salient example of how expression of some genes are often associated with a specific stressor, when in reality their expression is more likely a universal environmental stress response (ESR).

### *Cold challenge elicits a much stronger response than heat challenge in* A. poculata

Our data demonstrate that *A. poculata* exhibit greater behavioural and transcriptomic responses to the cold challenge applied here when compared to heat challenge, which is surprising considering that cold challenge temperatures were within *A. poculata’s* thermal range, while heat challenge temperatures were not (Figure 1B). In fact, the heat challenge exceeded any temperature experienced within their native environment over the last decade. Few studies have directly contrasted a coral’s response to thermal extremes in parallel, and studies that have demonstrated mixed results. In a tropical coral (*Acropora millepora*), Nielsen *et al*. (2020) observed improved coral condition under cold temperatures relative to ambient or heated conditions. In contrast to our results, heat stress causes a larger bleaching response than cold stress in *Aiptasia* (Bellis & Denver 2017). Conversely, Roth & Deheyn (2013) found that acute cold stress was more detrimental to the tropical coral *Acropora yongei* than heat stress, but did suggest that heat stress may be more detrimental over longer temporal scales. In a study investigating the responses of oysters *(Crassostrea gigas)* to heat and cold stress, Zhu *et al*. (2016) observed similar transcriptional responses to both stressors. While there is no clear consensus among studies, it is widely accepted that the specific temperatures reached in each stress treatment and the rate at which those temperatures are reached are both important factors (McLachlan *et al*. 2020). Heat challenge in our study may have elicited a more muted response, because *A. poculata* were collected in the summer, so these colonies were likely acclimated to warmer conditions, which would have made the cold challenge more stressful.

### Astrangia poculata *exhibits a convergent stress response repertoire to cold and heat challenge*

Despite highly divergent temperatures reached between temperature challenge experiments, we observed convergent behavioral and transcriptomic responses in *Astrangia poculata*. First, we observed reductions in feeding behaviour under both thermal challenges (Figure 2B), which were corroborated with convergent downregulation of genes associated with *locomotion* (GO: 0040011) and *response to mechanical stimulus* (GO: 0009612). *DELTA-thalatoxin-Avl2a* (AVL2A) was downregulated under both challenges; thalatoxin and other toxins are used while feeding in cnidarians (Schmidt *et al*. 2019) and are categorized under the *nematocyst* (GO:0042151) GO category. Furthermore, *myosin regulatory light polypeptide 9* (MYL9) was downregulated under both thermal challenges (Figure 4B) and this gene plays an important role in cell contractile activity via phosphorylation (Kumar *et al*. 1989) and may be instrumental for coral heterotrophy. Reduced polyp activity under thermal challenges may be due to temperatures exceeding local high and low temperatures or corals could be entering quiescent states, where lowered metabolic activity acts as an adaptation to extreme temperatures (Stuart & Brown 2006). Our transcriptomic results do not support quiescence and instead suggest large scale protein catabolism, which often occurs during starvation after an organism has metabolized most of its carbohydrate and lipid stores (Kaur & Debnath 2015; Davies *et al*. 2016). Increases in catabolic-related pathways point instead to high energetic demands associated with stress-related cell functions at both thermal extremes (Kültz 2005).

The other major convergent response observed under both thermal challenges was a generalized stress response. For example, *glutathione transferase activity* (GO: 0004364) was upregulated under both temperature extremes and this GO term is associated with detoxification of environmental pollutants and oxidative stress response in tropical corals (Downs *et al*. 2005). In addition, most enriched GO categories observed in both thermally-challenged *A. poculata* were involved in maintenance of the proteasome (Figure 5A-C). The role of the proteasome (discussed above) is integral to degradation and catabolism of oxidized proteins (Davies 2001) and may be important for the activation of NF-kB under stress (Rape & Jentsch 2002). These enriched GO terms across wide thermal challenges highlight conserved ESR pathways under both heat and cold thermal challenges.

In addition to convergently enriched GO terms, a number of individual genes were differentially expressed under both challenges. *Lon protease homolog 2, peroxisomal* (LONP2) was highly up-regulated in both experiments and this gene is involved in degradation of oxidatively damaged mitochondrial genes (Yang *et al*. 2018). LONP2 has been shown to be upregulated under high temperatures and under heavy metal stress in oysters (Sanni *et al* 2008). Additionally, *protein-glucosylgalactosylhydroxylysine glucosidase* (PGGHG) was downregulated under both thermal challenges (Figure 4B). PGGHG is a catalyst for the hydrolysis of glucose in hydroxylysine-linked residues of collagen (and collagen-like) proteins (Hamazaki & Hamazaki 2016). It is also a major component of isolated collagens from other marine invertebrates (e.g. sea anemones, Katzman *et al*. 1972), which are very reactive to a range of stressors (Traylor-Knowles 2019). Taken together, LONP2 and PGGHG play important roles in aposymbiotic *A. poculata’*s stress response.

The mitogen-activated protein kinase (MAPK) signaling pathway is key for mediating cell differentiation and apoptosis (Whitmarsh 2010) and has been previously implicated in a coral’s response to environmental stimuli (Sun *et al*. 2013). *A. poculata* consistently upregulated *increased sodium tolerance 1* (IST1) under both thermal challenges, which is also known as *putative MAPK-activating protein* (PM28; Figure 4B). In addition to IST1, *Tumour necrosis factor receptor 3* (TRAF3) was also highly upregulated under both thermal stressors (Figure 4B). TRAF3 is an intracellular signaling molecule that regulates MAPK activity and nuclear factor-*κ*B (Nf-*κ*B) signaling (Häcker 2011), which has been shown to be upregulated during stress-induced bleaching in *Aiptasia* (Mansfield *et al*. 2017). TRAF3 is constitutively upregulated or “front-loaded” in corals that are tolerant to heat stress (DeSalvo *et al*. 2010; Barshis *et al*. 2013; Seneca & Palumbi 2015) and is upregulated under low magnesium (Yuyama & Higuchi 2019), white band disease (Libro *et al*. 2013), and high carbon dioxide treatments (Kaniewska *et al*. 2012) in various coral species. Our results provide supporting evidence that TRAF3, along with IST1 and LONP2, may be part of the coral ESR and demonstrate consistent upregulation in response to various stressors, not just high temperatures.

## Conclusions

While stress response repertoires of tropical reef-building corals have been widely studied, especially in response to upper thermal extremes, this study represents the first to characterize the stress response of a naturally aposymbiotic coral to divergent thermal challenges. Our results demonstrate a strong response to cold challenge and a comparatively muted response to heat challenge. In addition, we provide evidence for a convergent stress response to divergent thermal challenges in *A. poculata* that is consistent with responses observed for tropical obligate coral species, which is surprising given the absence of symbiont-associated reactive oxygen species. The repertoire of convergent responses to thermal challenges highlighted here will provide the foundation for future research to investigate how symbiosis influences the coral stress response. We identified a number of genes that are differentially regulated under both thermal challenges, suggesting a universal stress response in a core set of CRGs. This work highlights the benefits to studying facultatively symbiotic corals to disentangle stress responses of the coral host from their algal symbionts, and future work leveraging this facultative relationship may lead to a stronger mechanistic understanding of why coral dysbiosis is increasing in frequency in corals worldwide.

## Acknowledgements

This work was made possible through the Boston University Marine Program with special thanks to Justin Scace for husbandry of *A. poculata*. The authors extend appreciation to K. Sharp, R. Rotjan, S. Grace and the annual Astrangia Workshop hosted by Roger Williams University and Southern Connecticut State University for fostering creative conversations and collaborations leading to this work. Also, we thank SJS Wuitchik and HE Aichelman for helpful edits and insights on this manuscript.

## Funding

This work was funded by Boston University’s start-up package to S.W.D and the Boston University Marine Program (BUMP).

## Data Availability

All sequences are available from the NCBI SRI under accession PRJNA595158. Code for all analyses are attached in supplementary materials, and are also available at https://github.com/wuitchik along with transcriptome files.

## Author Contributions

S.W.D designed the experiment. A.A., S.A.B., J.D.C., M.B.L., J.L.R., M.K.S., and I.F.T. conducted the experiment. B.E.B. and C.L.R. completed all molecular work and TagSeq library preparations. D.M.W. performed all statistical and bioinformatic analyses and drafted the manuscript. S.W.D. supervised the experiment, analyses and co-authored the manuscript. All authors edited and approved the manuscript.

## Notes

https://github.com/wuitchik/Divergent-thermal-challenges-elicit-convergent-stress-signatures-in-aposymbiotic-Astrangia-poculata

### Competing Interest Statement

The authors have declared no competing interest.

